# A comprehensive survey and benchmark of deep learning-based methods for atomic model building from cryo-EM density maps

**DOI:** 10.1101/2025.03.02.641021

**Authors:** Chenwei Zhang, Anne Condon, Khanh Dao Duc

## Abstract

Advancements in deep learning (DL) have recently led to new methods for automated construction of atomic models of proteins, from single-particle cryogenic electron microscopy (cryo-EM) density maps. We conduct a comprehensive survey of these methods, distinguishing between direct model building approaches that only use density maps, and indirect ones that integrate sequence-to-structure predictions from AlphaFold. To evaluate them with better precision, we refine standard existing metrics, and benchmark a subset of representative DLmethods against traditional physics-based approaches using 50 cryo-EM density maps at varying resolutions. Our findings demonstrate that overall, DL-based methods outperform traditional physics-based methods. Our benchmark also shows the benefit of integrating AlphaFold as it improved the completeness and accuracy of the model, although its dependency on available sequence information and limited training data may limit its usage.

## Introduction

Single-particle cryogenic electron microscopy (cryo-EM) is a major technique to discover the atomic structure of biomolecules at high resolution [1, 2]. This method relies on reconstructing a 3D electron density map from 2D images of frozen samples, and then fitting coordinates of atoms with it, a process termed *atomic model building*. As of 9 January 2025, more than 40000 electron microscopy density maps have been deposited to the Electron Microscopy Data Bank (EMDB) [3], with the number of released maps showing an exponential growth. Yet, only approximately 57 % of the associated atomic coordinates are resolved in the Research Collaboratory for Structural Bioinformatics Protein Data Bank (RCSB PDB) [4], as shown in Figure 1. As conventional methods for protein model building progressively refine an atomic model by minimizing a physics-based energy function (notably taking into account electrostatic interactions, steric clashes, and bond lengths), they indeed usually require substantial manual input to construct precise models [5–8]. To mitigate the need for extensive domain expertise and manual intervention, more automated tools and methods are needed.

**Figure 1:**
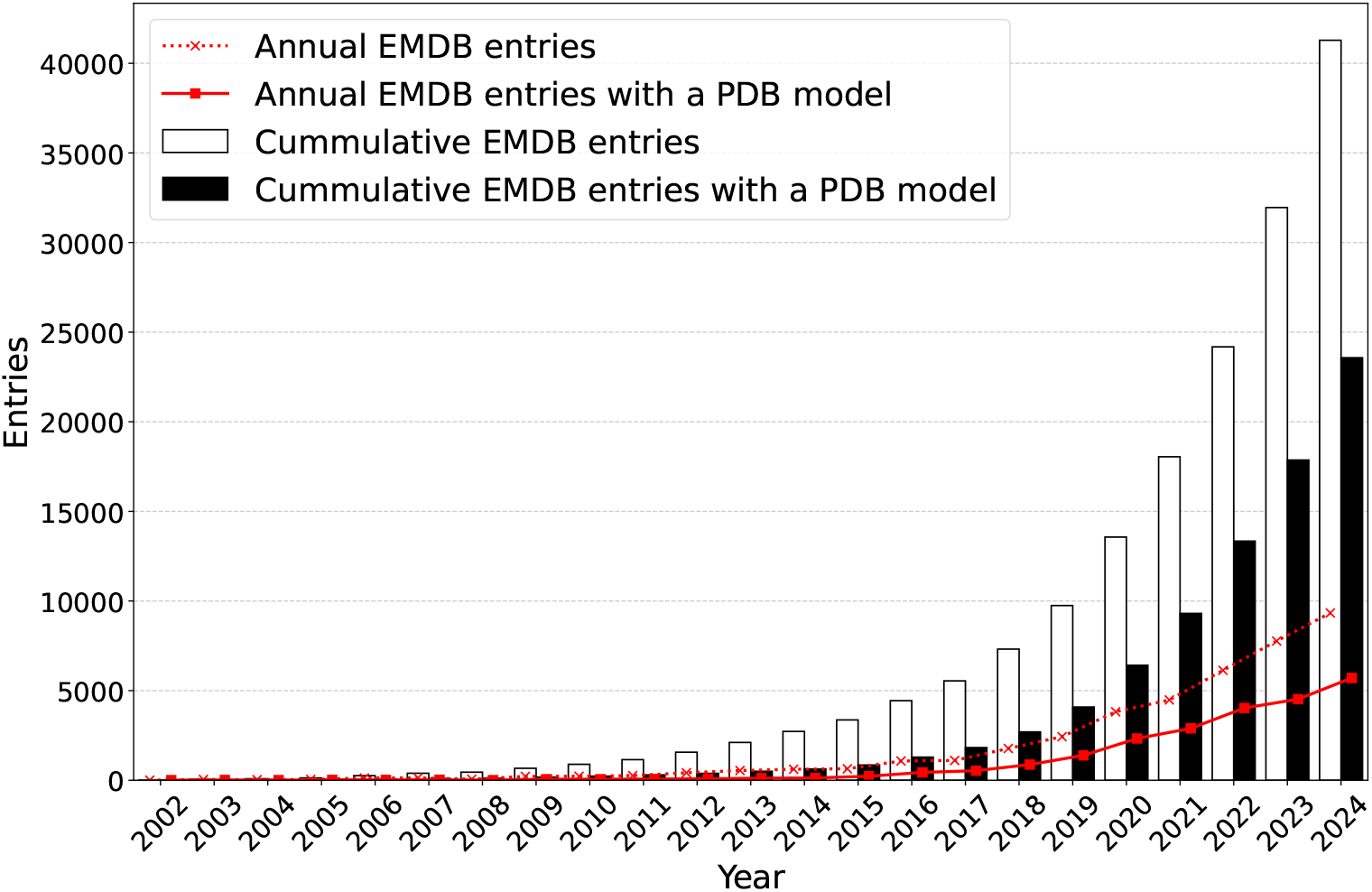
Bar chart: cumulative number of all EMDB entries and of those with an associated atomic structure in the PDB at the end of each year. Line plot: number of annually released EMDB and PDB entries. Data is shown until the end of 2024. The statistics were collected from EMDB on 2025-01-09 [3].

In this context, early data-driven methods that utilize classical machine learning (ML) techniques such as K-Nearest Neighbors (KNN) [9], K-means clustering [10], and Support Vector Machine (SVM) [11], have been proposed, but are limited to secondary structures or simplified backbone models, and not suited for large complex structures. More generally, automating model building is challenging, due to the presence of artifacts and noise, varying and inconsistent resolution caused by molecular flexibility or radiation damage, and the lack of homologous or predicted structures [5, 12, 13].

To address these limitations, various deep learning (DL)-based approaches have more recently been introduced, showing some significant advancements. Building upon previous reviews that have introduced physics-based, ML-based, and some DL-based methods for atomic model building from cryo-EM density maps [14–16], our review focuses on providing a timely and comprehensive survey of these recently proposed DL methods to better understand their common and distinctive features. Furthermore, we present a benchmark comparing several state-of-the-art methods, providing valuable insights and guidance for computer scientists, computational biologists and cryo-EM practitioners in the field. Our review is structured as follows:

1. We run a comprehensive survey of recent DL-based methods for automated model building, distinguishing methods that only use density maps, and indirect ones that integrate sequenceto-structure predictions from AlphaFold [17]. Among these two classes, we also cover the specific architectures, and the benefits and limitations of different methods.
2. To evaluate the performance of these methods, we motivate the need for and define new specific metrics to assess the correspondence between both global structural features and local features, that refine the standard Precision, recall, F1, and template matching (TM) scores.
3. Upon selecting four representative methods from (1) and using the metrics in (2), we evaluate and compare the performance of these methods by considering the alignment quality of predicted models from atomic to intermediate resolution, runtime and potential benefit of including structure prediction methods.

### Survey of DL algorithms for atomic model building

DL-based approaches initially focused on identifying secondary structure elements (SSEs) such as *α*-helix and *β*-sheets from density maps with resolution ranging from intermediate to low [18–21]. With the increasing availability of high-resolution density maps, recent DL techniques predict atom types and coordinates, from backbone structures that consist of a repeated sequence of nitrogen (N), C_*α*_, and beta-carbon (C_*β*_) atoms to full-atom models that include both backbone (main-chain) and side-chain atoms [13, 22–26]. Concurrently, the recent revolution in protein structure prediction led by AlphaFold and similar methods leveraging AI [17, 27–33], led to the integration of these sequence-to-structure approaches for atomic model building from density maps. In this review, we thus categorize DL-based model-building strategies into two types, which are summarized in Table 1, and detailed next.

**Table 1:**
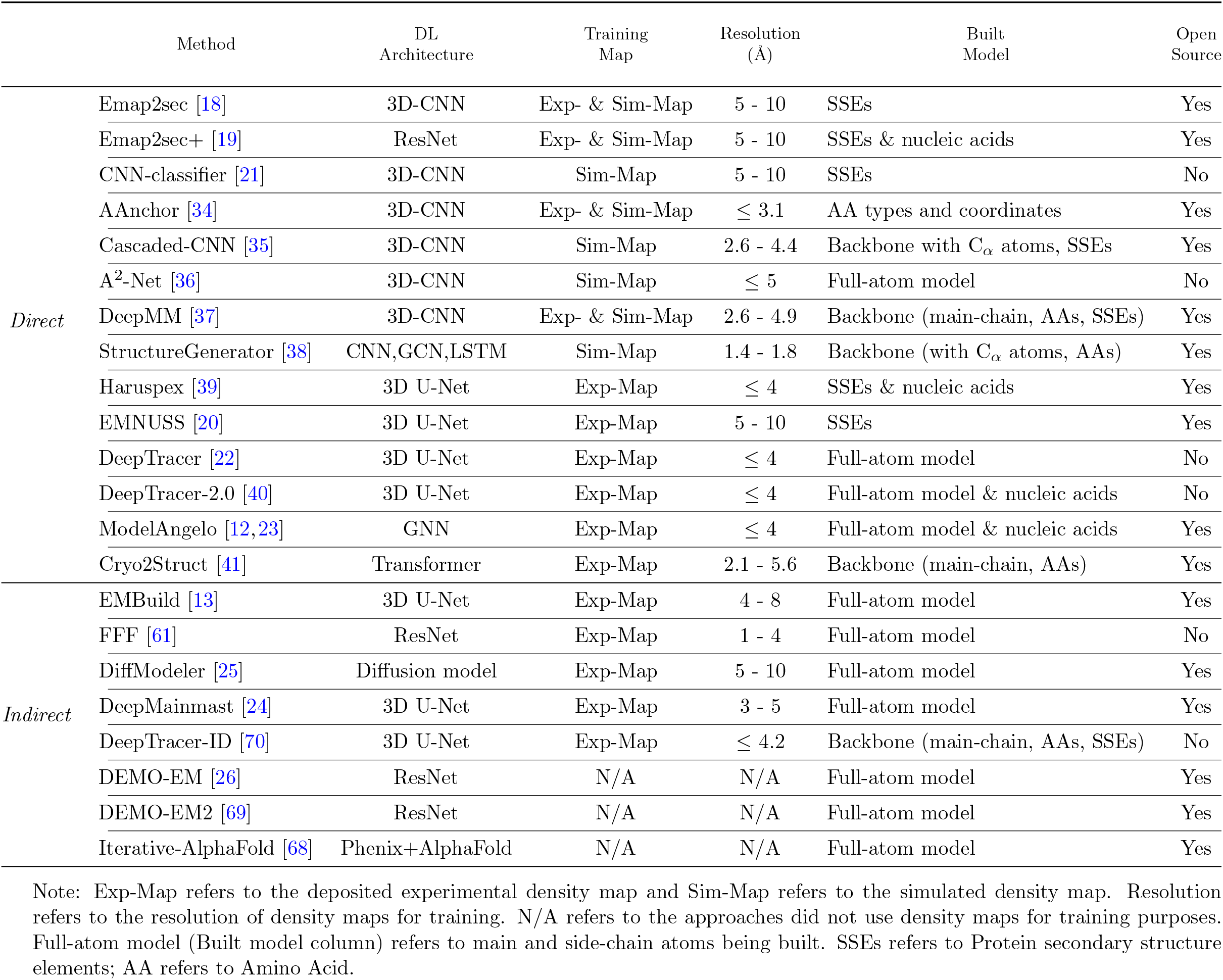
Summary of DL-based methods for atomic model building from cryo-EM density maps.

### Direct model building

Direct model-building methods [18–23, 34–41] leverage deep neural networks (DNNs) to predict voxel-wise representations from density maps directly, where each voxel is annotated with its associated backbone, SSE, amino acid type, C_*α*_ atom, and/or side-chain torsion angle. Backbone tracing, which connects predicted C_*α*_ atoms into chains, can be challenging due to disordered residues and map noise [35]. Advanced algorithms such as TSP solvers [42, 43], Vehicle Routing Problem (VRP) solvers [44, 45], and Monte Carlo Tree Search (MCTS) [46] are employed to address these challenges. Despite advancement, amino acid type prediction remains limited, mainly due to similarities among some amino acids within density maps. Improvements are achieved by aligning predicted amino acid sequences with target sequences and updating amino acid types accordingly [22, 23]. We illustrate this workflow in Figure 2a. Among direct model building methods, we can distinguish different types of DL architectures, as detailed next.

**Figure 2:**
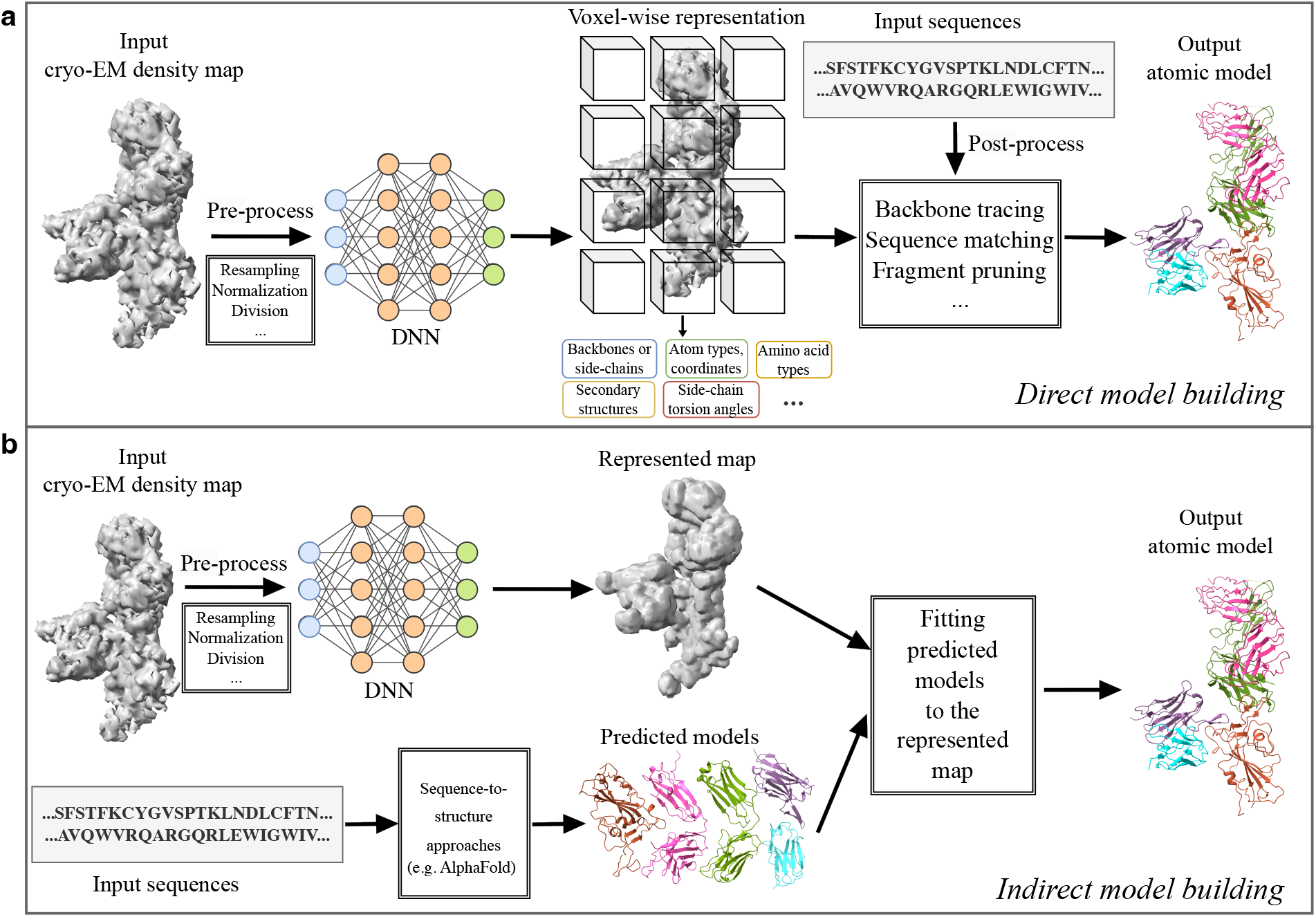
DL-based approaches (**a**) direct model building and (**b**) indirect model building. Note that the illustration of indirect model-building approaches represents an overview of selected methods. The illustrated protein complex structure is the SARS-CoV-2 spike in complex with antibodies B1-182.1 and A19-61.1 (PDB-ID: 7TBF; EMDB-ID: 25797; Resolution: 3.1 Å).

### CNN

Convolutional neural networks (CNNs) [47] excel in 2D image classification and have been extended to 3D density maps to predict protein structures, including SSEs as well as atom and amino acid types and coordinates. *Emap2sec* [18] adopts a two-phase 3D-CNN to identify SSEs from intermediate-resolution maps, improving accuracy by analyzing adjacent voxels, however it does not precisely locate SSEs coordinates. The improved version, *CNN-classifier* [21] refines SSE prediction, while it is limited to training on simulated maps. *AAnchor* [34] and *Cascaded-CNN* [35] specialize in annotating amino acids and constructing backbones respectively. Both are limited to high-resolution maps, and the latter lacks training on real-world maps. *A*^2^*-Net* [36] and *DeepMM* [37] focus on connecting and assigning predicted amino acids into individual chains to form complete protein structures, whereas they may overlook some structural features due to limitations inherent in their algorithms. *Structure Generator* [38] employs a pre-trained bidirectional long short-term memory (LSTM) network [48] to connect CNN-predicted amino acids and build protein models, but it is restricted to simulated maps. Despite these advancements, CNN-based methods struggle with rotational invariance [49, 50] which affects their performance on input density maps with varying orientations and in capturing global features and dependencies [51, 52] within density maps.

### U-Net

Originally designed for 2D medical image segmentation [53], U-Net architectures comprise down-sampling encoders and up-sampling decoders with skip connections to capture both the feature information and localization within the input map. *Haruspex* [39] leverages 3D U-Nets to identify protein SSEs and nucleic acids but sometimes misidentifies similar structures (e.g. confusing *β*-hairpin turns and polyproline helices with *α*-helical SSEs). On the other hand, *EMNUSS* [20] adopts an enhanced version of U-Net with extra connections (UNet++ [54]) to better identify *β*-sheets and coils. Yet, it still faces challenges with low-resolution maps and complex structures, and neither tool can fully build detailed atom models directly. Evolved from Cascaded-CNN, *Deep-Tracer* [22] utilizes multiple 3D U-Nets to identify and locate different protein structures and has shown promise in modeling coronavirus-related structures, despite occasional inaccuracies in C_*α*_ tracing and connectivity faults. *DeepTracer-2*.*0* [40] extends this approach to protein-DNA/RNA complexes, enhancing its utility in structure modelling. While U-Nets are good at extracting features, they struggle with low-resolution maps where critical features are less defined, and are prone to overfitting because of their complex architectures and limited high-resolution data available.

### GNN

Graph neural networks (GNNs) [55, 56] are particularly effective in modeling systems where relationships and dependencies are crucial. They work by treating elements like atoms or amino acids as nodes and their connections—such as bonds or interactions—as edges. This architecture makes GNNs ideal for building protein models. *ModelAngelo* [12, 23], a tool built on GNNs, enhances protein modeling by integrating cryo-EM map features with protein sequence embeddings derived from ESM-1b [57], a pre-trained protein large language model. However, the performance of GNN-based methods such as ModelAngelo is adversely affected when the node-edge relationships in low-resolution maps are ill-defined.

### Transformer

The Transformer architecture [58], initially a breakthrough in natural language processing, has also been effective in analyzing 2D images [59] and 3D volumes [60]. Attributable to its multi-head attention mechanism, Transformers can emphasize important areas in density maps, such as SSEs, improving the accuracy of protein model construction. *Cryo2Struct* [41] employs this technique to identify protein atoms and amino acids by learning the interactions between atoms across long distances within the map. A major limitation of Transformers is their large number of parameters, making them prone to overfitting, particularly when trained on limited or highly specific datasets like cryo-EM density maps. Additionally, hyperparameter tuning in Transformers is non-trivial and can greatly impact model performance.

### Indirect model building

Some indirect model-building methods [13,25,61] utilize DNNs to enhance the structural features of original maps, yielding representations like backbone probability maps [61], main chain probability maps [13], or traced backbone maps [25]. These methods differ from direct model-building approaches since they rely on sequence-to-structure predictions like AlphaFold to construct models of protein chains, domains, or complexes. These predicted models are then aligned to the represented maps using various fitting algorithms, including rigid-body [62, 63], partial [64], semi-flexible [13], and flexible [65–67] fitting, to assemble a complete model. We illustrate this workflow in Figure 2b. In addition, some methods [24] integrate structures predicted by AlphaFold into the C_*α*_ tracing stage to improve the model construction. Other methods [26, 68, 69] leverage map features to refine preliminary structures predicted by approaches like AlphaFold. In contrast, some strategies [70] utilize AlphaFold to refine the initial models derived directly from density maps. Similarly to direct model building methods, we can distinguish the following architectures.

### U-Net-Based

*EMBuild* [13] uses a deep U-Net architecture trained on intermediate-resolution density maps to generates main-chain probability (MCP) maps, which extract volumetric information of main-chain atoms. The AlphaFold-predicted chain models are then aligned to MCP maps and assembled to complete protein structures. Nevertheless, the performance of EMBuild may decline if the fitting is sub-optimal. *DeepMainmast* [24] employs deep U-Nets to identify amino acids, atom types, and C_*α*_ positions in density maps. It improves C_*α*_ tracing by integrating AlphaFold predictions and aligning them to the map. The final model is selected from the best direct builds or AlphaFold fits. DeepMainmast also has a version called *DeepMainmast-Base*, which builds models directly from maps. Nonetheless, both approaches demand significant computational hours due to intensive optimizations. *DeepTracer-ID* [70] begins with a preliminary atomic model derived from DeepTracer and iteratively refines it by leveraging AlphaFold predictions. However, this approach is suitable only for high-resolution maps and protein sequences longer than 100 amino acids.

### ResNet

Residual networks (ResNets) [71], a variant of CNNs, use residual connections that allow the layers to learn from inputs directly, helping to prevent gradient vanishing during training. The residual connection allows for the use of much deeper layers, improving predictions in deeper neural networks designed for complex data. For instance, *FFF* [61] adapts a variation of ResNet, known as RetinaNet [72], to generate backbone probability maps and pseudo-peptide vectors, which facilitate fitting Alphafold predictions into these maps, while this method is sensitive to map resolution. *DEMO-EM* [26] leverages a deep ResNet to predict distances between protein domains, assisting in the flexible assembly of atomic models. Since these domain structures are produced without map data, however, they may not accurately reflect the actual protein structure. Built upon DEMO-EM, *DEMO-EM2* [69] intertwines both chain-level and domain-level fittings to enhance model accuracy. Although deeper ResNet models can perform better, they increase the complexity of networks and can be challenging to optimize.

### Diffusion model

Diffusion models [73] are used to reduce noise from structured data, such as images or audios. Extended to 3D density maps, *DiffModeler* [25] takes advantage of denoising diffusion implicit models [74] to learn protein backbone features from intermediate-resolution density maps. Using these learned representations, DiffModeler generates traced backbone maps that are used to fit AlphaFold-predicted single-chain models into a full-atom model. A caveat is that diffusion models require considerable computational resources for both training and inference.

### Iterative-AlphaFold

Terwilliger *et al*. [68] have introduced an iterative procedure for Phenix that refines AlphaFold predictions by implicitly incorporating cryo-EM density maps (*Iterative-AlphaFold*). This process involves tweaking an initial AlphaFold model to better fit the map and then using the adjusted model as a custom template for next-cycle AlphaFold predictions. This procedure enhances the model’s accuracy but works best with high-resolution maps due to rebuilding constraints inherent in Phenix [75].

### Evaluation metrics for comparison of predicted and target models

Evaluation metrics for model building depend on the alignment of predicted and target models, where the alignment of C_*α*_ atoms accounts for the alignment of residues. Given a predicted structure containing *L*_pred_ residues and a corresponding target structure with *L*_target_ residues, the number of aligned C_*α*_ atoms (i.e. paired atoms), denoted as *L*_align_, follows the relationship: *L*_align_ *≤ L*_pred_ *≤ L*_target_. Existing metrics such as recall, precision, F1-score, and sequence recall [23], C_*α*_ matching score [5], C_*α*_ quality score [41], and sequence recall [23] are commonly used to measure the percentage of paired C_*α*_ atoms. However, these metrics count paired atoms without considering chain correspondence. Ideally, paired atoms should reside within the same corresponding chains. As illustrated in Figure 3a, C_*α*_ atoms *A* from the target structure and *A*^*′*^ from the Phenix-predicted structure belong to different corresponding chains: *A* in the yellow chain, while *A*^*′*^ in the blue chain. Metrics that disregard chain correspondence incorrectly pair *A* with *A*^*′*^, artificially inflating the precision, recall, and F1-score. In contrast, when chain correspondence is considered, *A* and *A*^*′*^ should not be paired since they belong to different corresponding chains, resulting in significantly lower TM-score [76]. In addition, although the TM-score accounts for chain correspondence, it does not consider the accuracy of residue pairing identity, i.e., whether the paired residues share the same amino acid type, as illustrated in Figure 3b. These limitations highlight the need to design better metrics.

**Figure 3:**
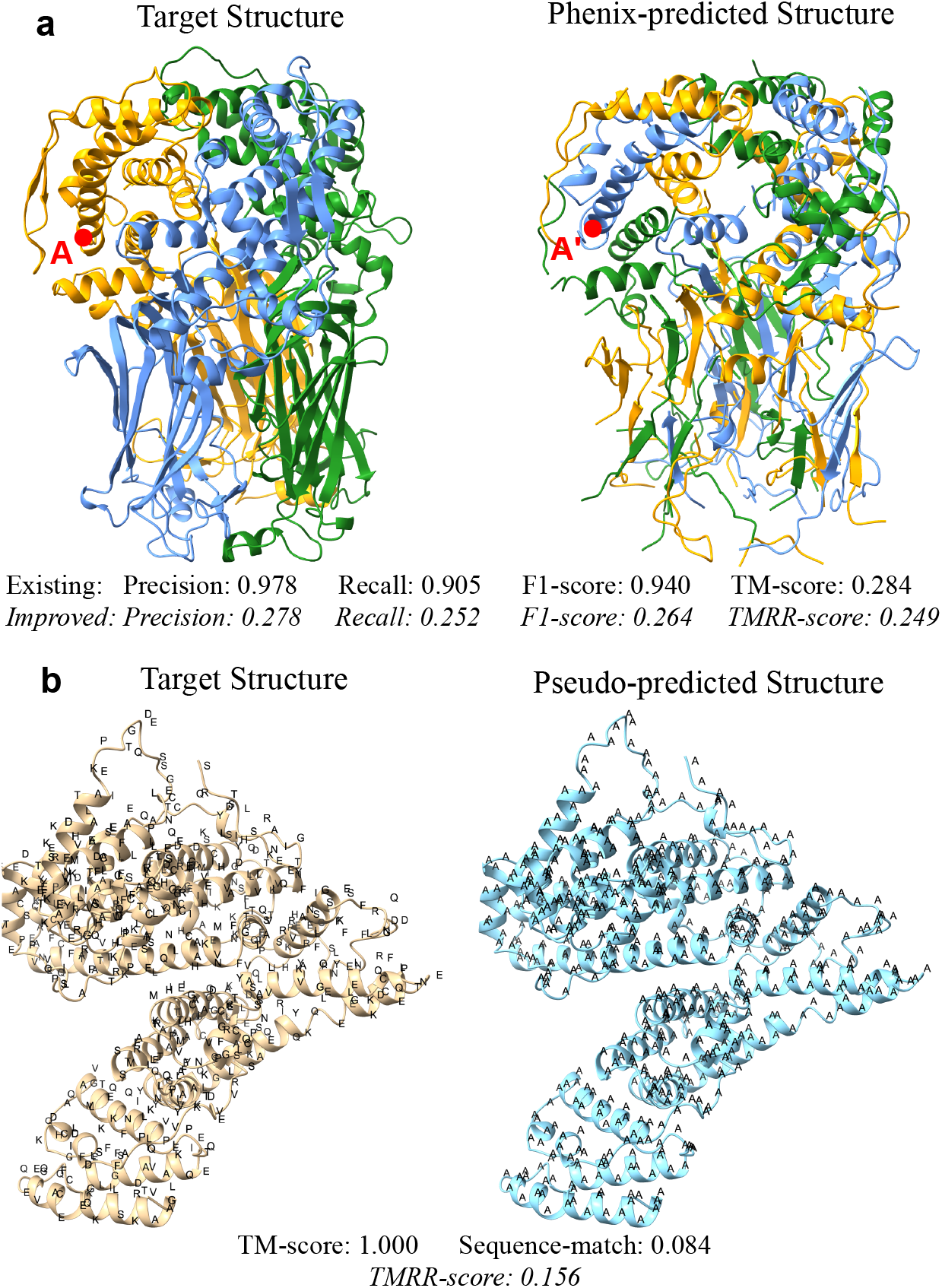
Comparison of prediction scores using different evaluation metrics. **(a)** The overall topology of the Phenix-predicted protein structure closely resembles that of the target one. However, the folding patterns of the predicted protein chains are distinctly different. Existing metrics, such as those outlined by ModelAngelo [23], do not account for chain correspondence, thereby resulting in very high precision, recall, and F1-score. Conversely, the TM-score, which considers chain correspondence, is notably low. In our improved metrics, these scores are aligned, showing minimal discrepancies. The visualized protein is a structure of rotavirus VP6 (PDB-ID: 3J9S). Different chains are depicted in different colors. The red dots *A* and *A*^*′*^ refer to C_*α*_ atoms in the target and Phenix-predicted structures, respectively. **(b)** The pseudo-predicted structure shares an identical shape with the target one, achieving a TM-score of 1.000. However, they consist of distinct amino acid types, resulting in a low Sequence-match score of 0.084. In our improved metric, TMRR-score, this value is 0.156. The visualized protein is a structure of human Alpha-fetoprotein (PDB-ID: 7YIM). The letter represents the one-letter code for amino acids.

### Improved evaluation metrics

As discussed above, the limitations of existing evaluation metrics include (i) the ignorance of chain-level correspondence, and (ii) the absence of residue identity matching in TM-score. To address (i), we refine the precision, recall and F1-score, by using a heuristic algorithm, US-align [77], to systematically align the predicted and target structures based on the highest TM-score obtained from all possible combinations of their chains. To address (ii), we introduce a new metric, named ***TMRR-score***, that couples the TM-score with Residue-recall to simultaneously measure both structural and residue-type similarities.

### Refined precision, recall, and F1-score

First, we align the two structures using US-algin and collect the aligned C_*α*_ atom of each residue from one structure and its counterpart from the other. We define a residue as paired, or a C_*α*_ atom as correctly predicted, if the distance between the aligned C_*α*_ atoms is less than or equal to a specified threshold of 3 Å [5]. Next, we leverage the defined correctly-predicted C_*α*_ atom, termed CP-C_*α*_ atom, to calculate the following quantitative measures.

*Precision* is defined as the percentage of predicted C_*α*_ atoms that are correct, expressed as:

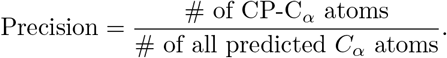

*Recall* is defined as the fraction of C_*α*_ atoms in the target structure that are correctly identified, expressed as:

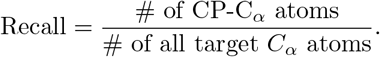

*F1-score* is the harmonic mean of precision and recall, expressed as:

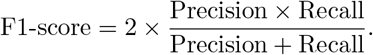

### TMRR-score

Inspired by the harmonic mean [78] that is more sensitive to the lower value, we define the *TMRR-score* by coupling the TM-score and Residue-recall:

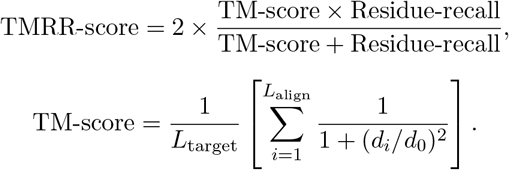

Where 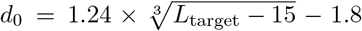 is a length-dependent scale to normalize the C_*α*_ atom distance of the *i*th pair of aligned residues, *d*_*i*_ [76]. *Residue-recall* is defined as the ratio of paired residues that match in amino acid type to the overall count of residues within the target structure.

The TMRR-score ranges from (0, 1] with a higher score indicating a more pronounced similarity in the context of bot’ shape and residue identity. A score of 1 indicates that predicted and target models are identical, i.e. *L*_align_ = *L*_pred_ = *L*_target_. Note that we design Residue-recall to measure the accuracy of amino acid type predictions, unlike Sequence-match, which simply measures the percentage of correctly predicted amino acids relative to the total number of predicted residues. Residue-recall accounts for residue coverage. For instance, a predicted structure covering only a fraction of the target structure could still achieve a high Sequence-match score. We suggest that the sensitivity of the TMRR-score to lower values helps mitigate the risk that an exceptionally high score in one metric could compensate a poor score in the other, thus encouraging a more balanced and unbiased evaluation.

### Benchmarking results

To our knowledge, previous work has not conducted a comprehensive comparison across cutting-edge model-building methods, and the evaluation metrics used vary and are not unified across different methods. Using our new evaluation metrics, we conducted a comprehensive benchmarking of four cutting-edge model-building approaches. Phenix serves as the baseline, and the others are DL-based, namely ModelAngelo [12], EMBuild [13], and DeepMainmast [24]. We selected these three DL-based methods because they are fully automated, open-source, and have consistently demonstrated state-of-the-art performance based on thorough assessments. ModelAngelo is a direct model-building method, whereas EMBuild operates indirectly. DeepMainmast offers two modes: one that integrates AlphaFold predictions (direct) and another does not (indirect). Specifically, we benchmarked these methods over a diverse set of 50 proteins including monomers, homomers, and heteromers, with their density maps at resolution ranging from 1.8 Å to 7.8 Å (as detailed in Supplementary Table S1). Note that these 50 candidate maps exclude maps from the training datasets of the DL approaches to ensure an unbiased and fair comparison. For more details about our implementation, see SI text.

### Performance comparison across methods

In Figure 4, each box-and-whisker plot illustrates the distribution of scores for atomic models constructed from 50 density maps, each assessed using the respective methods^1^. Note that for the DeepMainmast and DeepMainmast-Base results, we opted to benchmark the constructed backbone models instead of the full-atom models, since we observed significant distortions in some full-atom models derived from their backbone models using Rosetta-v2021.16, as illustrated in Supplementary Figure S1. Given that our evaluation metrics specifically focus on C_*α*_ atoms, they primarily assess the accuracy of backbones, ensuring that the metrics are not biased towards either backbone-only or full-atom models.

**Figure 4:**
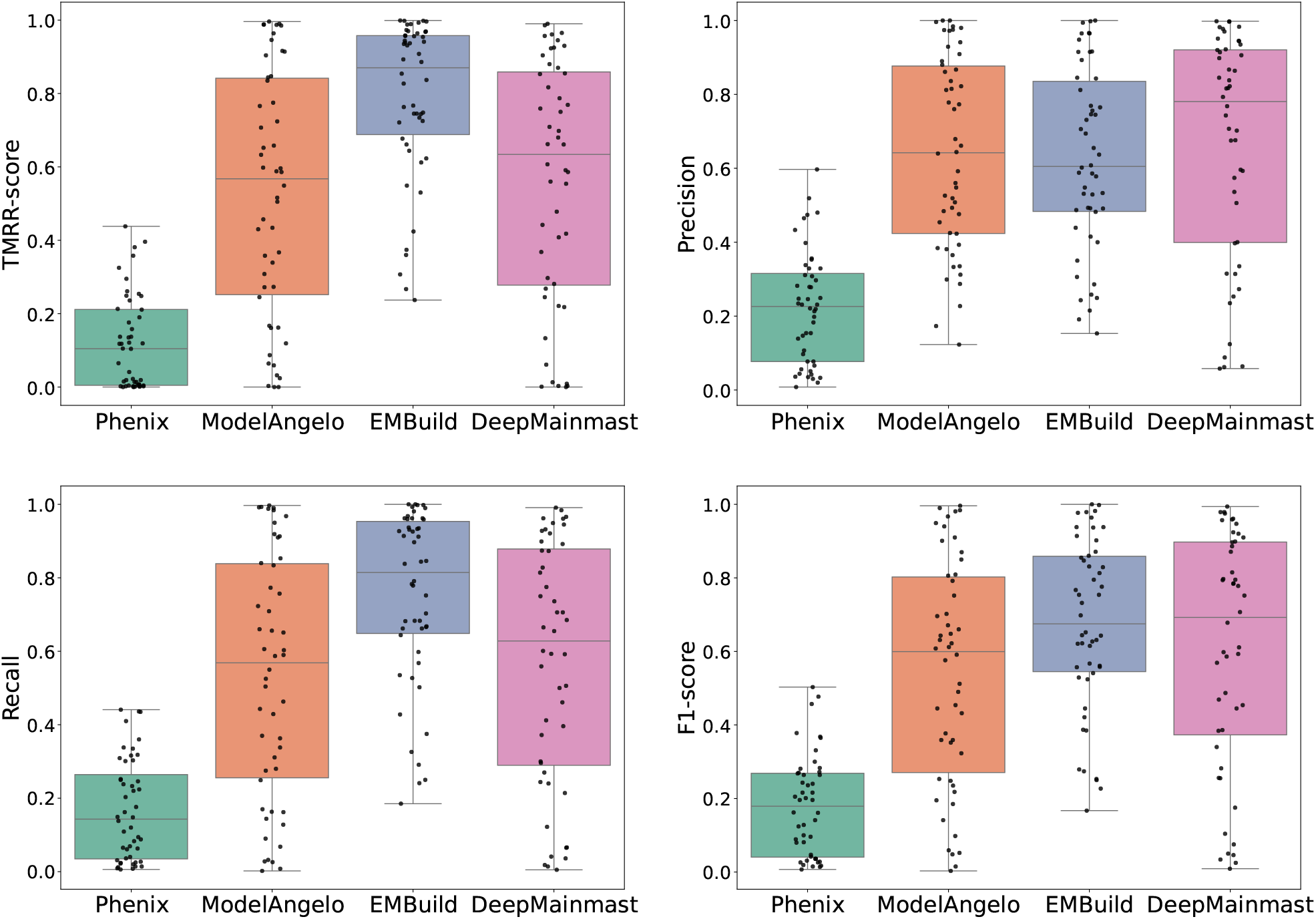
Comparison of improved metrics (TMRR-score, Precision, Recall, and F1-score) of 50 protein models generated by Phenix, ModelAngelo, EMBuild, and DeepMainmast. The dots represent the scores for each protein model.

The TMRR-scores revealed that EMBuild outperformed its counterparts, exhibiting exceptional model reliability. Moreover, EMBuild showed the narrowest box width (interquartile range), indicating its more consistent results across the tested density maps. Both ModelAngelo and DeepMainmast achieved comparable mean and median TMRR-scores above 0.5, suggesting that their constructed models closely match the target ones. In contrast, the lower scores of Phenix reflected its underperformance in accurately building protein models and predicting amino acid types. Additionally, DeepMainmast showed the highest mean and median precision, suggesting its effectiveness in producing relevant structures. On the other hand, EMBuild scored the highest mean values in both recall and F1-score, with the narrowest box width emphasizing its high modeling coverage and overall performance, aligning with its TMRR-score. Table 2 summarizes the mean and median scores for each method. Based on these results, we concluded that DL-based methods involving EMBuild, ModelAngelo, and DeepMainmast all showcased good performance across varying evaluation metrics, demonstrating their capabilities to build accurate atomic models from density maps. We also employed existing metrics including TM-score [76], precision, recall, and F1-score [23] to compare each method, with the results presented in Supplementary Figure S2. We observed significant discrepancies and inconsistencies between TM-scores and the other three measures. For instance, Phenix exhibited a very low median TM-score below 0.2, while achieving relatively high scores in the other three metrics. These findings reinforce the necessity of using our metrics.

**Table 2:**
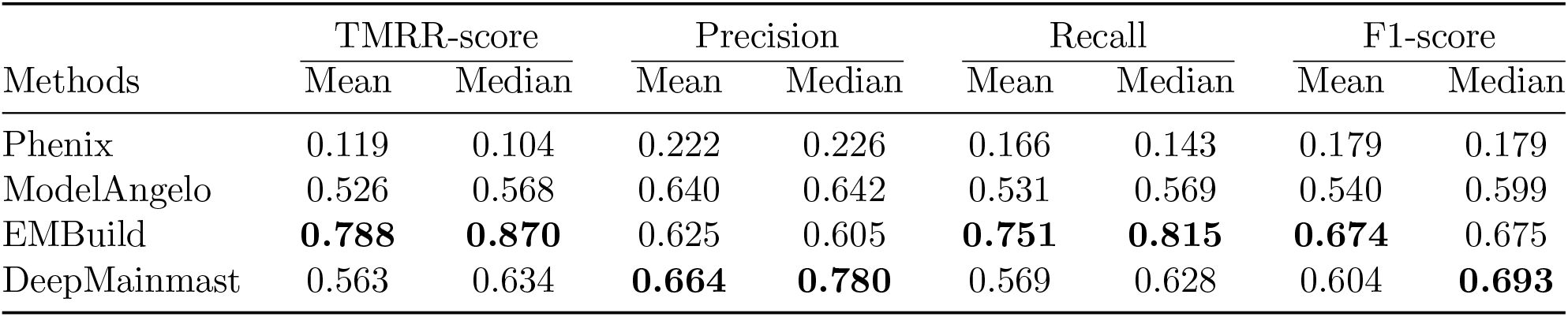
Performance comparison of different model-building methods using improved metrics.

### Performance comparison across map resolutions

We compared different methods against density maps of varying resolutions from 1.8 to 7.8 Å, and their TMRR-scores were depicted as a heat map in Figure 5. ModelAngelo demonstrated superior performance at near-atomic resolutions, consistently achieving scores above 0.5, indicating its robustness in constructing meaningful protein models. However, its performance declined at resolutions worse than 3.7 Å. DeepMainmast also performed well at high resolutions, although some models were poorly constructed. In contrast, EMBuild consistently exhibited good performance across a broad range of resolutions, suggesting its independence from map resolution. Phenix, on the other hand, was highly dependent on resolution, with TMRR-scores plummeting to nearly zero at resolutions worse than 4 Å. Furthermore, we observed that all methods except EMBuild struggled to build meaningful models from intermediate-resolution density maps (4 8 Å). Interestingly, DeepMainmast was ineffective at very high resolutions (1.8 - 2.5 Å), suggesting that its algorithms failed to adequately capture map features at these finer resolutions. Additional heat maps of precision, recall, and F1-score are displayed in Supplementary Figures S3-S5, and the results align with their TMRR-scores.

**Figure 5:**
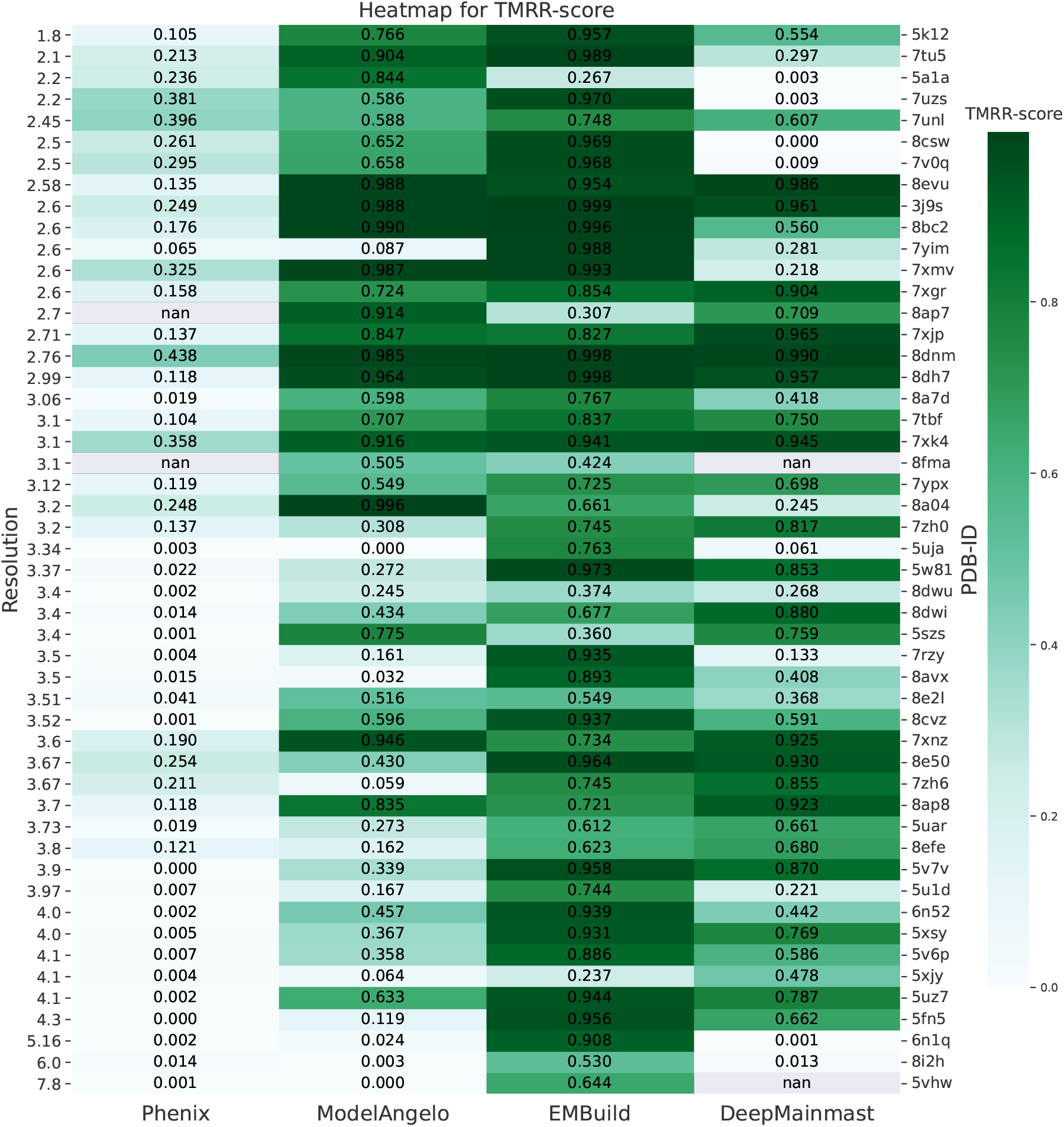
The heat map for the TMRR-score of 50 protein models generated by Phenix, ModelAngelo, EMBuild, and DeepMainmast, sorted by their map resolution from high to low. The darker color refers to the higher score. The value in each cell represents the specific score for the generated model. The left labels show the resolutions and the right labels show the PDB IDs of each generated model.

### Visual comparison

While the performance comparison in the previous section provides valuable insights from an overall comparison on a large dataset, further insights can be gained by visually examining specific cases where each method performs poorly. To this end, we visualized the constructed atomic models from three representative high-resolution density maps. Figure 6a shows a poorly ModelAngelo-constructed model with a lower TMRR-score than that of others. We observed that some regions that were enclosed by black boxes were not constructed by ModelAngelo, causing the incompleteness and low TMRR-score, although the rest parts were well constructed. These incompletely modeled areas can be attributed to regions with high noise and low-density values. Figure 6b shows a poorly constructed model by EMBuild. We found that some parts of the model were misaligned with the corresponding density volumes, thereby causing an inferior construction. Moreover, we noticed that the completeness was high in the EMBuild-constructed model and almost every density volume was registered due to its explicit usage of AlphaFold predictions. Figure 6c reveals a poorly constructed model by DeepMainmast, while perfectly constructed by the other two methods with TMRR-scores above 0.9. Although DeepMainmast also integrated AlphaFold in their model-building process, rather than directly aligning the AlphaFold predictions to the map as EMBuild did, it leveraged the predictions to enhance the C_*α*_ tracing process. Therefore, DeepMainmast’s performance was still strongly dependent on the map quality.

**Figure 6:**
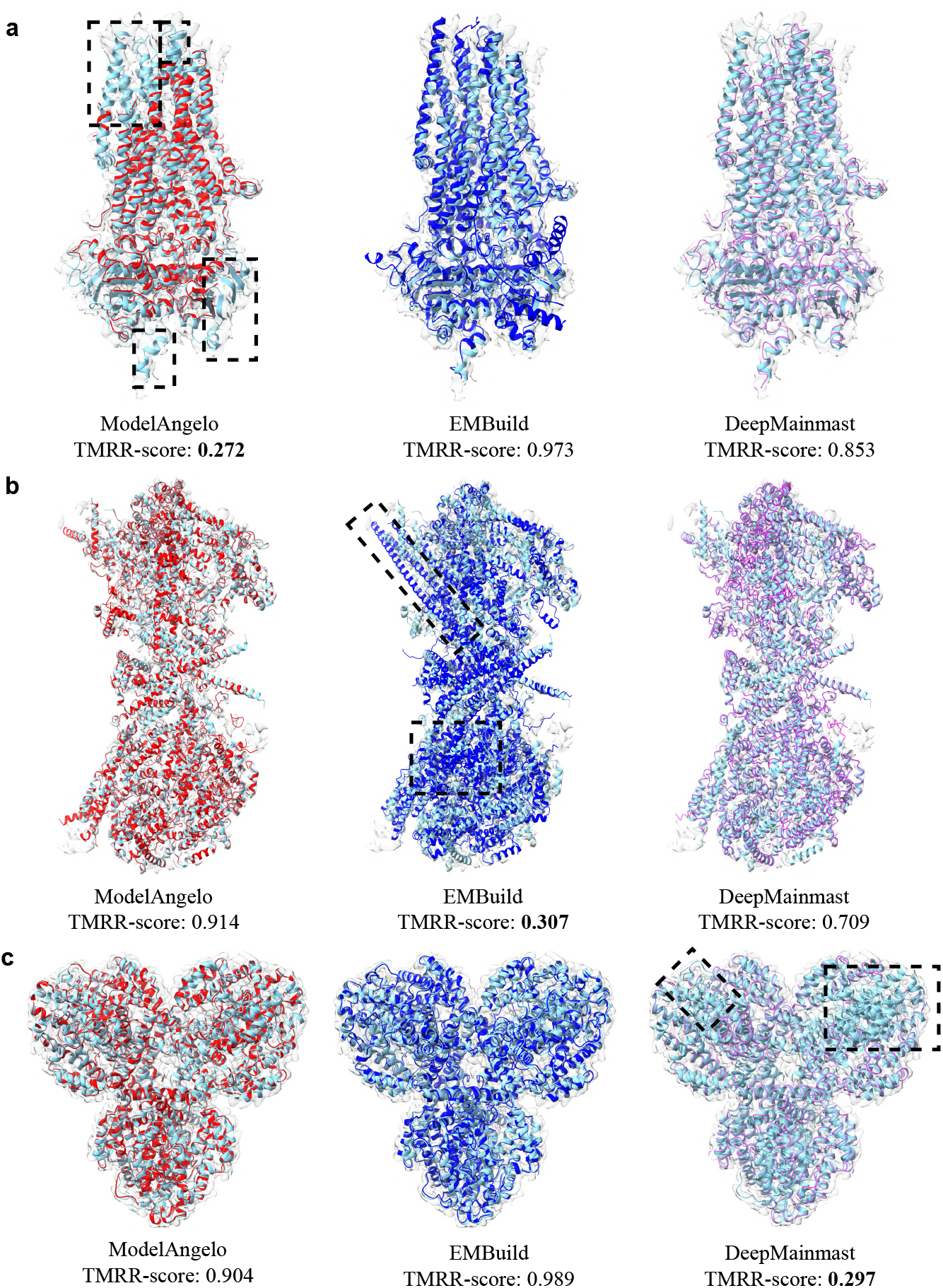
**a-c** Constructed protein atomic models by ModelAngelo (red), EMBuild (blue), and DeepMainmast (purple), respectively, with cryo-EM density maps (transparent gray) and the corresponding reference PDB structures (cyan). Each TMRR-score value is shown at the bottom of the model. Black boxes highlight the poorly modeled regions. **(a)** Phosphorylated, ATP-bound structure of zebrafish cystic fibrosis transmembrane conductance regulator (PDB-ID: 5W81; EMDB-ID: 8782; Resolution: 3.37 Å). **(b)** Membrane region of the Trypanosoma brucei mitochondrial ATP synthase dimer (PDB-ID: 8AP7; EMDB-ID: 15560; Resolution: 2.7 Å). **(c)** Structure of the L. blandensis dGTPase in the apo form (PDB-ID: 7TU5; EMDB-ID: 26126; Resolution: 2.1 Å).

### Impact of incorporating AlphaFold

To investigate how AlphaFold enhances model-building performance, we compared DeepMainmast, which incorporated AlphaFold, against DeepMainmast-Base, which did not. Figure 7a shows that DeepMainmast outperformed DeepMainmast-Base across all four metrics and produced more consistent results, as evidenced by its narrower box width. Additionally, we visually compared the models constructed by both methods for the same map, as illustrated in Figure 7b. The results revealed that DeepMainmast can build more complete and less fragmented models than DeepMainmast-Base, because of the integration of AlphaFold. Both quantitative and visual results suggest that integrating AlphaFold enhances modeling coverage, achieving a higher degree of completion and reducing fragmentation and disconnections. This achievement stems from AlphaFold’s proficiency in constructing a complete topology. However, integrating AlphaFold comes at a cost: increased computational demands and extended runtime. As evidenced in Table 3, DeepMainmast has a significantly slower runtime compared to DeepMainmast-Base.

**Table 3:** Model building time taken by various methods across different sequence lengths. All methods were implemented on one NVIDIA A100 GPU with AMD EPYC 7V12 64-Core Processor.

| Sample | Sequence length | Model building runtime (hours) |  |  |  |  |
| --- | --- | --- | --- | --- | --- | --- |
|  |  | Phenix | ModelAngelo | EMBuild | DeepMainmast | DeepMainmast-Base |
| EMDB-27755 | 495 | **0.29** | 1.31 | 2.30 | 6.07 | 3.34 |
| EMDB-28081 | 1053 | **0.65** | 1.32 | 3.29 | 9.82 | 3.73 |
| EMDB-27574 | 2288 | 1.86 | 2.38 | **1.80** | 42.66 | 13.54 |
| EMDB-27022 | 4648 | **1.43** | 3.99 | 3.82 | 41.31 | 25.52 |

**Figure 7:**
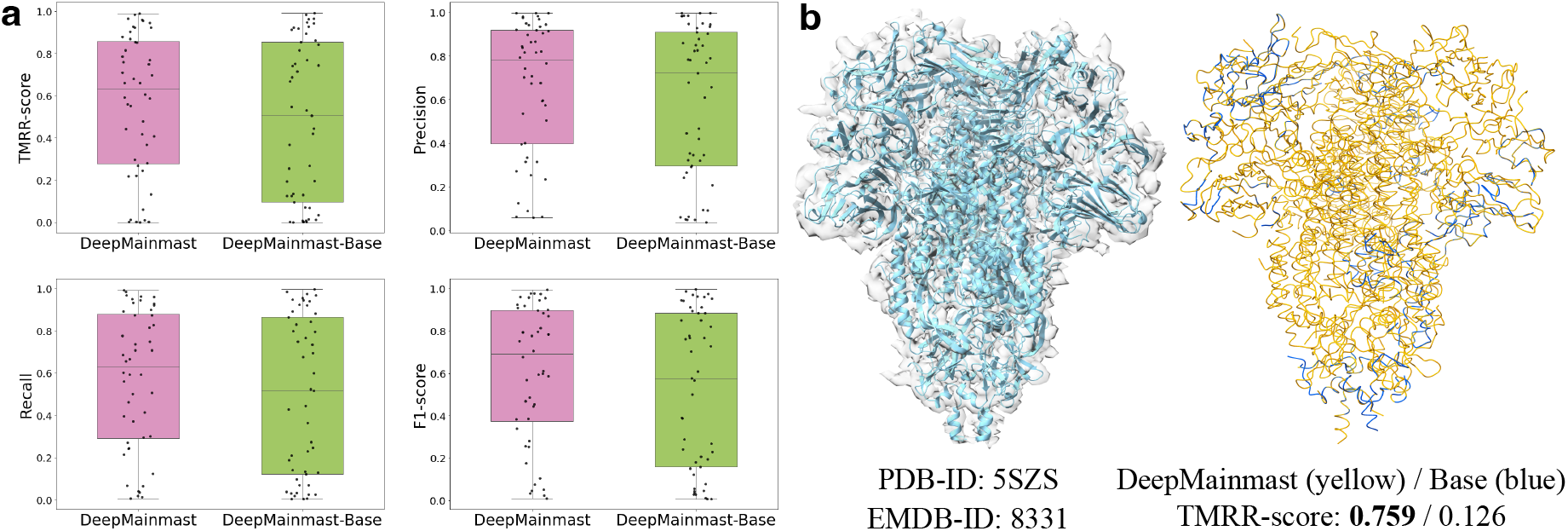
**(a)** Comparison of four metrics (TMRR-score, Precision, Recall, and F1-score) of 50 protein models generated by DeepMainmast and DeepMainmast-Base. The dots represent the scores for each protein model. **(b)** Left: the reference PDB structure colored in cyan superimposes onto the corresponding cryo-EM density map colored in transparent gray. Right: the superimposed constructed atomic models by DeepMainmast and DeepMainmast-Base, colored in yellow and blue, respectively. TMRR-score values are shown on the bottom of the models. Glycan shield and epitope masking of a coronavirus spike protein (PDB-ID: 5SZS; EMDB-ID: 8331; Resolution: 3.4 Å).

### Runtime comparison

The runtime for Phenix included map sharpening and model-building processes; for ModelAngelo included the initial model-building step and three-round refinements; for EMBuild included AlphaFold prediction and model-building process; and for DeepMainmast and DeepMainmast-Base included model-building process and AlphaFold prediction for the former. Note that the AlphaFold prediction runtime refers to the time to run AlphaFold locally, as some predictions were not available in the AlphaFold `Protein Structure Database`. Table 3 lists the runtime comparison across various methods for constructing atomic models with different sequence lengths, ranging from 495 to 4648 residues. There is a noticeable variance in computational efficiency, from minutes to days. Phenix demonstrated the fastest runtime for both short and long sequences. ModelAngelo and EMBuild exhibited acceptable runtime, with a slight increase as the sequence length extended. In contrast, DeepMainmast and DeepMainmast-Base showed considerable runtime, with the runtime increasing exponentially as the sequence length extended. Nevertheless, all methods were significantly faster than manual model building, which requires extensive domain expertise.

## Conclusion and future directions

Deep learning-based approaches have significantly advanced automated atomic model building from cryo-EM density maps. In this review, we present a comprehensive survey of recent state-of-the-art approaches, categorizing them into direct methods that rely solely on density maps and indirect methods that incorporate sequence-to-structure predictions from tools like AlphaFold. We delve into the specific neural network architectures employed by these methods, examining their design and functionality. Furthermore, we discuss the pros and cons of each approach, providing insights into their effectiveness across various modeling scenarios.

To generate unbiased comparison and consistent results, we improved the existing evaluation metrics. Upon comparing the performance of four representative methods across 50 density maps spanning a variety of resolutions, our results demonstrate that DL-based methods including ModelAngelo, EMBuild, and DeepMainmast significantly outperform the physics-based approach embodied by Phenix. EMBuild stands out across the entire spectrum of tested resolutions and excels in terms of TMRR-score, recall, and F1-score, showcasing its capability in constructing reliable and accurate atomic models, with the caveat that occasionally EMBuild completely misses the mark. On the other hand, ModelAngelo and DeepMainmast also perform well, particularly at high resolutions, with DeepMainmast achieving the highest precision.

The integration of AlphaFold into the model-building process has significantly enhanced construction completeness and accuracy. This improvement is particularly evident in the performance of EMBuild and DeepMainmast, where the inclusion of AlphaFold predictions boosts modeling coverage (completeness). However, AlphaFold’s dependency on available sequence information limits its applicability. Thus, alternative approaches such as ModelAngelo remain crucial for constructing atomic models in the absence of sequence data.

The field of automated atomic model building from cryo-EM density maps is still in its early stage of development and stands to benefit notably from enhanced collaboration between machine learning scientists and structural biologists. The rapid emergence of deep learning architectures and algorithms that achieved significant success in other fields highlights the potential for more sophisticated DL-based approaches to improve atomic model building. Moreover, there is a need for methods that leverage multi-modal data, such as structural templates, amino acid sequences, and 2D cryo-EM projected images, for training purposes. Incorporating physico-chemical constraints, such as rotation angles and bond lengths, should also be considered in future network design. While most methods currently focus on high-resolution density maps, there is also a pressing need to develop methods suitable for low-resolution maps.

Finally, a critical barrier for the effective development of DL-based methods for automated atomic model building is the need for high-quality data—specifically, perfect map-model pairs. A large number of maps are poorly aligned with their corresponding models, hindering neural networks’ ability to learn effectively. In the future, concerted efforts from research institutions and pharmaceutical companies to share cryo-EM density maps and atomic models with the community would significantly enhance the diversity of current datasets, leading to reduced bias during network training and boost in performance.

## Supporting information

Benchmark Results

## Supplementary Data

Supplementary data are available in the attached files.

## Acknowledgments

This work is supported in parts by a NFRFE-2019-00486 grant.

## Data Availability

Our improved evaluation metrics are accessible at https://github.com/chenwei-zhang/cryoEVAL. We compiled a comprehensive list of papers detailing de novo atomic model building approaches with their source codes, which is accessible at https://github.com/chenwei-zhang/papers-model-building-from-cryoem-maps.

