## Supplementary Information for "A comprehensive survey and benchmark of deep learning-based methods for atomic model building from cryo-EM density maps"

The Supplemental Information contains:

- Supplementary text
- Supplementary Figures S1–S4
- Supplementary Table S1
- Supplementary File

### Benchmarking Implementation

We benchmarked the following software versions: Phenix-v1.20, ModelAngelo-v1.0.12, EMBuild (as of February 15, 2024), and DeepMainmast (as of March 13, 2024). Rosetta-v2021.16 was used to assist DeepMainmast in producing full-atom models. All evaluated DL-based approaches used pre-trained networks provided by the developers. The AlphaFold predictions used for both EMBuild and DeepMainmast were obtained from the `AlphaFold Protein Structure Database` [1]. For those predictions that were not available in the database, we generated them locally using AlphaFold-v2.3.2. All methods were implemented on one NVIDIA A100 GPU with AMD EPYC 7V12 64-Core Processor. Visualization of cryo-EM density maps and atomic models was produced by UCSF ChimeraX [2].

#### Supplementary Figures

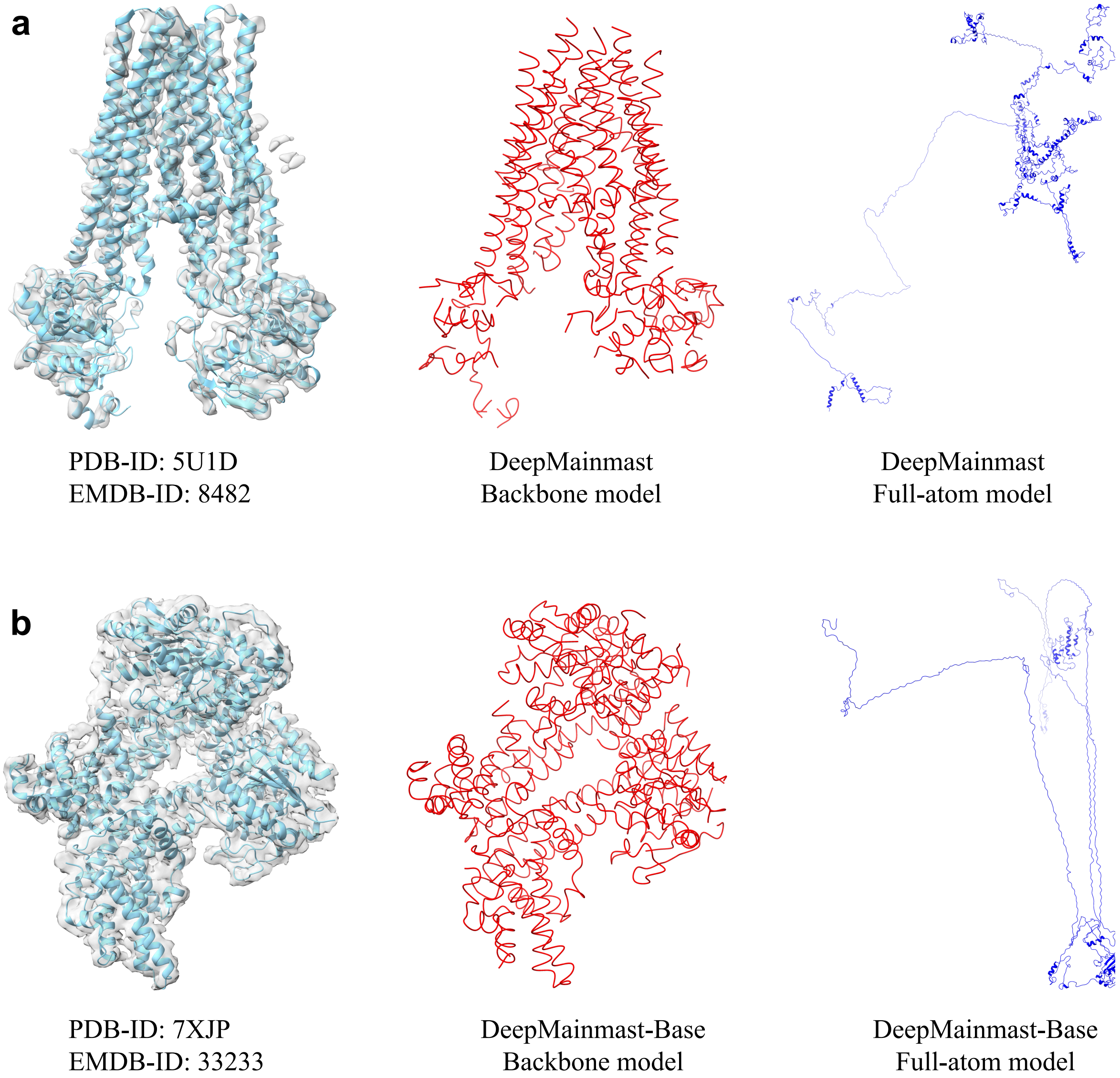


**Figure S1.** Left: the reference PDB structures colored in cyan superimposes onto the corresponding cryo-EM density maps colored in transparent gray. Middle: backbone models constructed by DeepMainmast (at **a**) and DeepMainmast-Base (at **b**), colored in red. Right: distorted full-atom models constructed by DeepMainmast (at **a**) and DeepMainmast-Base (at **b**), colored in blue. **(a)** The human TAP ATP-Binding Cassette Transporter (PDB-ID: 5U1D; EMDB-ID: 8482; Resolution: 3.97 Å). **(b)** EDS1 and SAG101 with ATP-APDR (PDB-ID: 7XJP; EMDB-ID: 33233; Resolution: 2.71 Å).

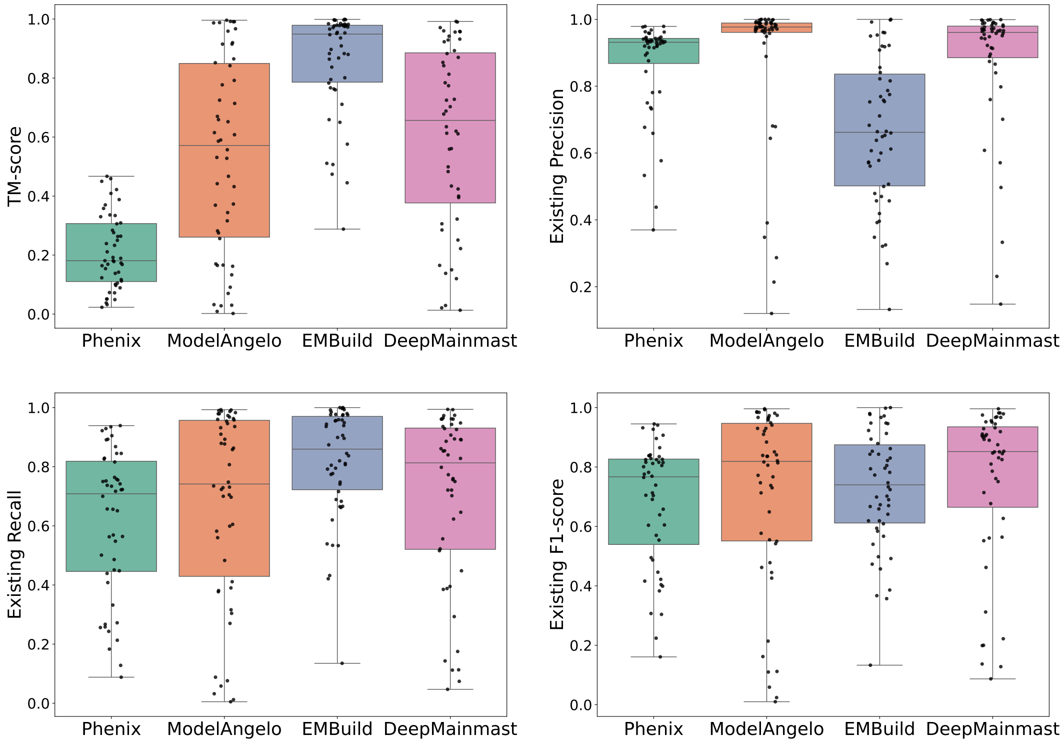


**Figure S2.** Comparison of existing metrics (TM-score, Existing Precision, Existing Recall, and Existing F1-score) of 50 protein models generated by Phenix, ModelAngelo, EMBuild, and DeepMainmast. The dots represent the scores for each protein model.

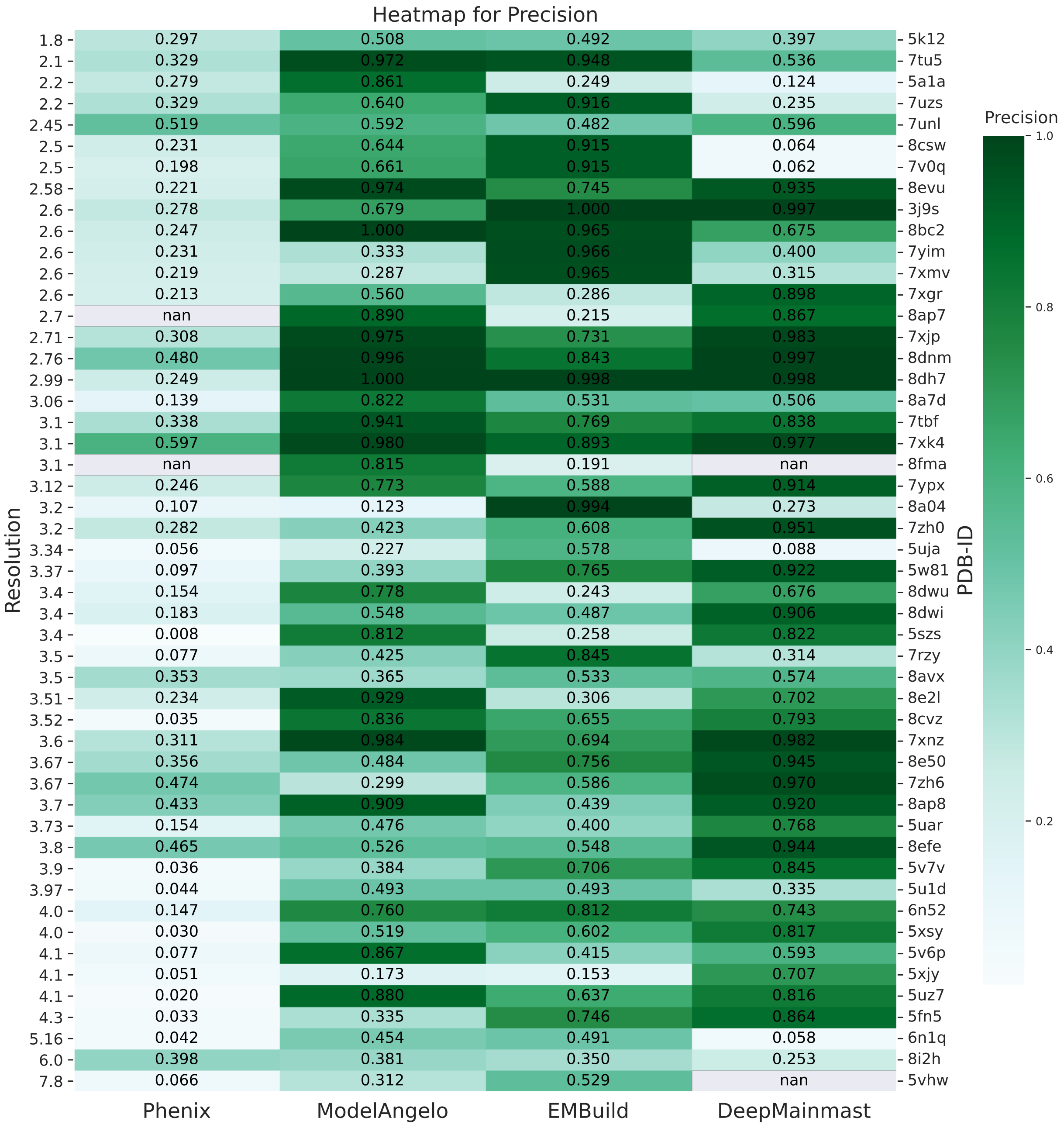


**Figure S3.** The heat map for precision of 50 protein models generated by Phenix, ModelAngelo, EMBuild, and DeepMainmast, sorted by their map resolution from high to low. The darker color refers to the higher score. The value in each cell represents the specific score for the generated model. The left labels shows the resolutions and the right labels shows the PDB IDs of each generated model.

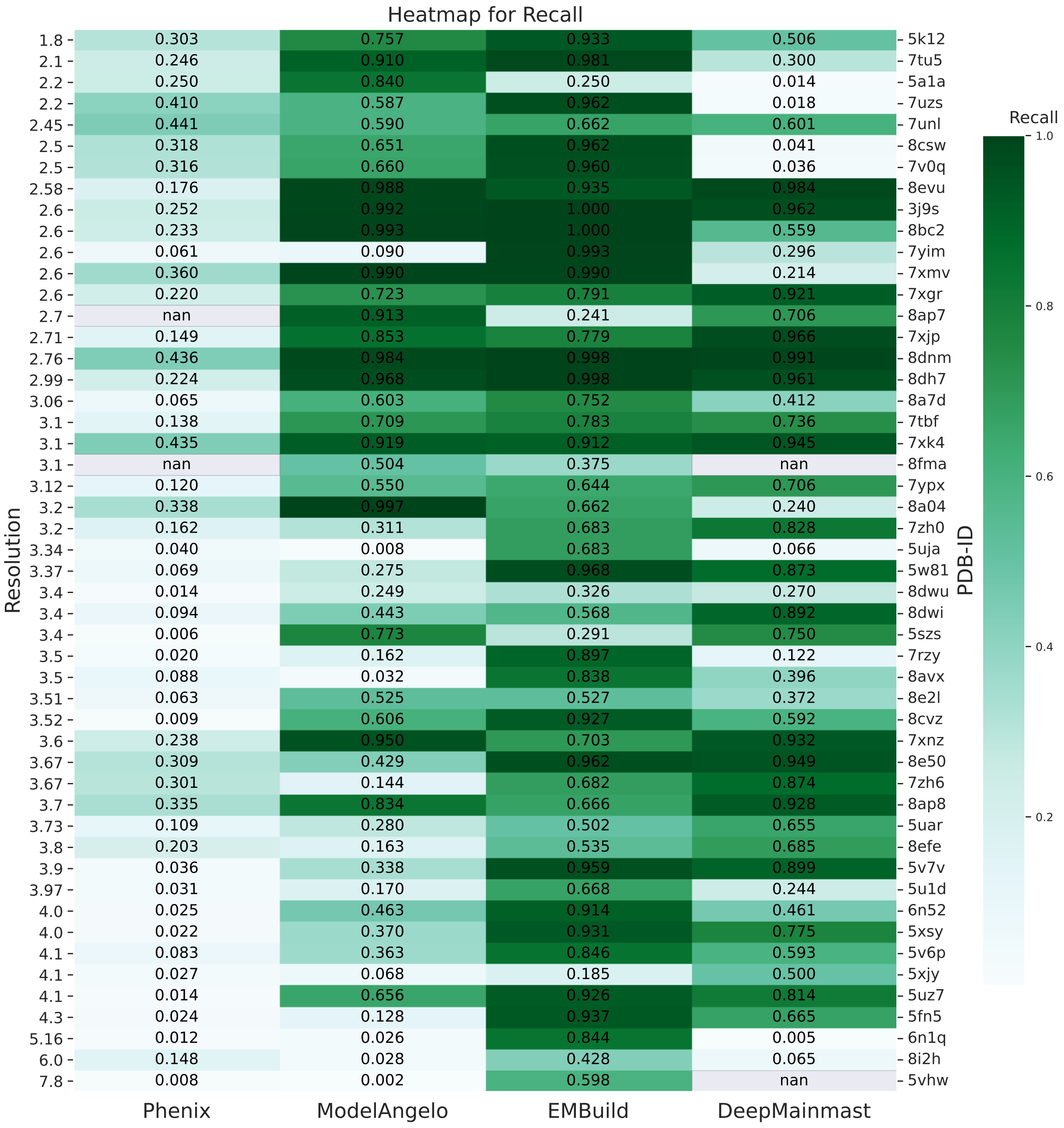


**Figure S4.** The heat map for recall of 50 protein models generated by Phenix, ModelAngelo, EMBuild, and DeepMainmast, sorted by their map resolution from high to low. The darker color refers to the higher score. The value in each cell represents the specific score for the generated model. The left labels shows the resolutions and the right labels shows the PDB IDs of each generated model.

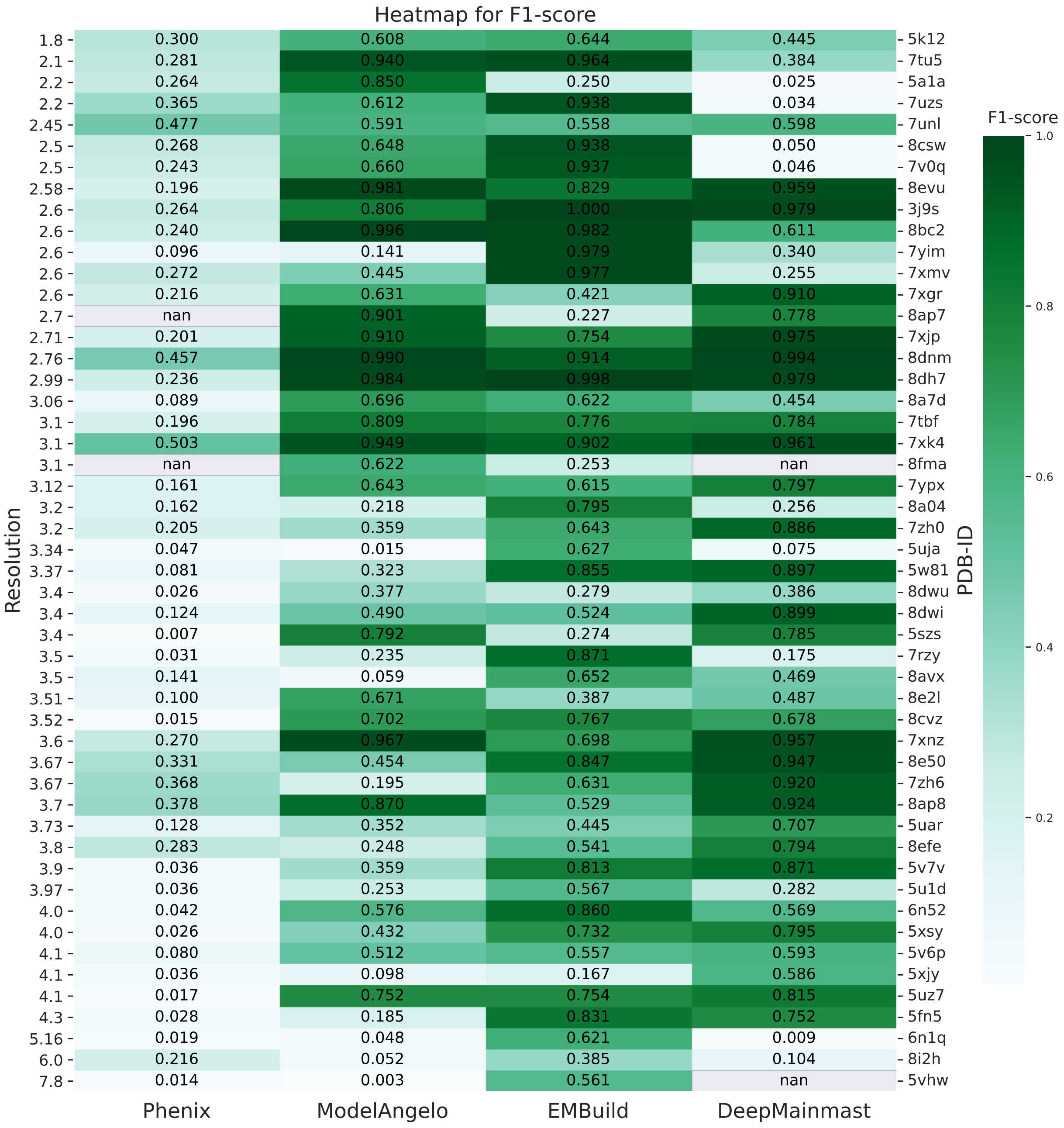


**Figure S5.** The heat map for F1-score of 50 protein models generated by Phenix, ModelAngelo, EMBuild, and DeepMainmast, sorted by their map resolution from high to low. The darker color refers to the higher score. The value in each cell represents the specific score for the generated model. The left labels shows the resolutions and the right labels shows the PDB IDs of each generated model.

#### Supplementary Table

**Table S1.** Summary of benchmarking data.

| EMID | PDB ID | Resolution (Å) | # of Residues | Stoichiometry |
|---|---|---|---|---|
| 26917 | 7UZS | 2.2 | 691 | Monomer - A1 |
| 26948 | 7V0Q | 2.5 | 691 | Monomer - A1 |
| 33187 | 7XGR | 2.6 | 671 | Monomer - A1 |
| 33233 | 7XJP | 2.71 | 1160 | Hetero 2-mer - A1B1 |
| 33243 | 7XK4 | 3.1 | 1940 | Hetero 6-mer - A1B1C1D1E1F |
| 33306 | 7XMV | 2.6 | 1926 | Homo 6-mer - A6 |
| 33331 | 7XNZ | 3.6 | 1912 | Homo 4-mer - A4 |
| 33861 | 7YIM | 2.6 | 609 | Monomer - A1 |
| 34017 | 7YPX | 3.12 | 1353 | Hetero 6-mer - A3B3 |
| 14716 | 7ZH0 | 3.2 | 556 | Monomer - A1 |
| 14725 | 7ZH6 | 3.67 | 556 | Monomer - A1 |
| 15047 | 8A04 | 3.2 | 717 | Homo 3-mer - A3 |
| 15220 | 8A7D | 3.06 | 3239 | Hetero 3-mer - A1B1C1 |
| 15560 | 8AP7 | 2.7 | 4822 | Hetero 30-mer - A2B2C2D2E2F2G2H2I2J2K2L2M2N2O2 |
| 15561 | 8AP8 | 3.7 | 1165 | Hetero 5-mer - A1B1C1D1E1 |
| 15686 | 8AVX | 3.5 | 1876 | Homo 2-mer - A2 |
| 15960 | 8BC2 | 2.6 | 2180 | Homo 10-mer - A10 |
| 26973 | 8CSW | 2.5 | 691 | Monomer - A1 |
| 27022 | 8CVZ | 3.52 | 4648 | Hetero 10-mer - A4B4C2 |
| 27431 | 8DH7 | 2.99 | 1052 | Homo 2-mer - A2 |
| 27574 | 8DNM | 2.76 | 2288 | Homo 4-mer - A4 |
| 27755 | 8DWI | 3.4 | 495 | Monomer - A1 |
| 27760 | 8DWU | 3.4 | 3366 | Hetero 9-mer - A5B3C1 |
| 27899 | 8E50 | 3.67 | 767 | Monomer - A1 |
| 28081 | 8EFE | 3.8 | 1053 | Hetero 2-mer - A1B1 |
| 28637 | 8EVU | 2.58 | 1934 | Hetero 6-mer - A2B1C1D1E1 |
| 29290 | 8FMA | 3.1 | 22242 | Hetero 22-mer - A11B11 |
| 27842 | 8E2L | 3.51 | 3267 | Hetero 6-mer - A4B2 |
| 8637 | 5V6P | 4.1 | 814 | Homo 2-mer - A2 |
| 26626 | 7UNL | 2.45 | 1008 | Homo 2-mer - A2 |
| 6272 | 3J9S | 2.6 | 1194 | Homo 3-mer - A3 |
| 8331 | 5SZS | 3.4 | 3975 | Homo 3-mer - A3 |
| 2984 | 5A1A | 2.2 | 4088 | Homo 4-mer - A4 |
| 8194 | 5K12 | 1.8 | 3348 | Homo 6-mer - A6 |
| 26126 | 7TU5 | 2.1 | 2784 | Homo 6-mer - A6 |
| 24783 | 7RZY | 3.5 | 2310 | Homo 7-mer - A7 |
| 6724 | 5XJY | 4.1 | 2305 | Monomer - A1 |
| 0346 | 6N52 | 4 | 1742 | Homo 2-mer - A2 |
| 25797 | 7TBF | 3.1 | 873 | Hetero 5-mer - A1B1C1D1E1 |
| 6770 | 5XSY | 4 | 2029 | Hetero 2-mer - A1B1 |
| 8482 | 5U1D | 3.97 | 1522 | Hetero 3-mer - A1B1C1 |
| 3240 | 5FN5 | 4.3 | 1542 | Hetero 4-mer - A1B1C1D1 |
| 8623 | 5UZ7 | 4.1 | 1438 | Hetero 5-mer - A1B1C1D1E1 |
| 8461 | 5UAR | 3.73 | 1494 | Monomer - A1 |
| 8560 | 5UJA | 3.34 | 1460 | Monomer - A1 |
| 8642 | 5V7V | 3.9 | 818 | Monomer - A1 |
| 8782 | 5W81 | 3.37 | 1494 | Monomer - A1 |
| 9317 | 6N1Q | 5.16 | 4088 | Homo 8-mer - A8 |
| 35136 | 8I2H | 6 | 682 | Monomer - A1 |
| 8685 | 5VHW | 7.8 | 4228 | Homo 4-mer - A4 |

#### Supplementary File

All benchmarking results can be accessed at the Supplementary File:
***SI_all_benchmarking_results.xlsx***.
